# Genomic surveillance reveals circulation of multiple variants and lineages of SARS-CoV-2 during COVID-19 pandemic in Indian city of Bengaluru

**DOI:** 10.1101/2023.03.14.532352

**Authors:** Darshan Sreenivas, Vinay Kumar, Kandasamy Kathirvel, Rakesh Netha Vadnala, Sachin Mishra, Bhagyashree Shelar, Srilatha Marate, Lakshminarayanan CP, Sai Disha K, Manisha Bhardwaj, Awadhesh Pandit, Satyajit Mayor, Uma Ramakrishnan, Dimple Notani

## Abstract

Genomic surveillance in response to coronavirus disease (COVID-19) pandemic is crucial for tracking spread, identify variants of concern (VoCs) and understand the evolution of its etiological agent, severe acute respiratory syndrome coronavirus-2 (SARS-CoV-2). India has experienced three waves of COVID-19 cases, which includes a deadly wave of COVID-19 that was driven by the Delta lineages (second/Delta wave) followed by another wave driven by the Omicron lineages (third/Omicron wave). These waves were particularly dramatic in the metropolitan cities due to high population density. We evaluated the prevalence, and mutational spectrum of SARS-CoV-2 variants/lineages in one such megapolis, Bengaluru city, across these three waves between October 2020 and June 2022. 15,134 SARS-CoV-2 samples were subjected to whole genome sequencing (WGS). Phylogenetic analysis revealed, SARS-CoV-2 variants in Bengaluru city belonged to 18 clades and 196 distinct lineages. As expected, the Delta lineages were the most dominant lineages during the second wave of COVID-19. The Omicron lineage BA.2 and its sublineages accounted for most of the COVID-19 cases in the third wave. Most number of amino acid changes were observed in spike protein. Among the 18 clades, majority of the mutations and least similarity at nucleotide sequence level with the reference genome were observed in Omicron clades.

## Introduction

SARS-CoV-2 has spread throughout the world and claimed millions of lives. Once unknown etiological agent of unexplained Pneumonia in China towards the end of 2019 was found to be a novel Coronavirus of genus Beta by metagenomics studies^1^. The first sequence made available in January 2020 was crucial for COVID-19 diagnostics, genomic surveillance to track the spread of the virus and vaccine development. Further, it aided in designing of tiled multiplex primer-based amplicon sequencing and thereby high throughput SARS-CoV-2 genomic surveillance workflows^2,3^.

Base incorporation errors and/or along with selection pressure from intra host environment may favor emergence of new variants that can spread more easily, escape host immune responses and change clinical presentation. Since the world is globally connected by a variety of modes of fast transport, a novel virus or a new variant of a virus can spread over long distances and some of these have serious consequences on public health. Such SARS-CoV-2 variants are called variants of concern (VoCs)^4–6^. To combat this, various public health policymakers have advocated monitoring the importation of such variants from international travelers and containing their spread in the community by enforcing social distancing norms, lockdowns and contact tracing along with active genomic surveillance of COVID-19 cases in the community. However, once VoCs are seen in the community, a surge in COVID-19 cases is most likely^7,8^.

India saw its first COVID-19 case towards the end of January 2020; however, significant number of cases and the first death in the country from the state of Karnataka were reported only in March 2020. Since then, the country observed multiple epidemic waves and lockdowns^9^. The second and third epidemic waves correlate with dominant representation of Delta (B.1.617.2 and descendant lineages) and Omicron (B.1.1.529 and descendent lineages), respectively as circulating lineages. Despite experiencing a deadly wave of COVID-19 cases that was supposedly driven by the Delta lineages, robust genomic surveillance efforts to understand various lineages and their temporal dynamics before and during the Delta wave were not available from India, until the third wave. This is evident in increased number of sequences shared on Global Initiative on Sharing All Influenza Data (GISAID) during the third wave as a result of efforts by various laboratories across the country and the genomic surveillance consortiums, such as Indian SARS-CoV-2 genomics consortium (INSACOG). As a part of one such consortium (anchored by CSIR-CCMB) with participating institutes from multiple cities of India, we embarked on a hyperlocal sequencing effort in four major cities of India, New Delhi, Bengaluru, Pune and Hyderabad.

In this study, we sequenced 15,134 samples which had tested positive for SARS-CoV-2 RNA presence by Reverse Transcription Real-time polymerase chain reaction in Bengaluru, a city with a population of around 13.1 million. 11,159 sequences passed our QC for coverage >50% against the reference sequence (genbank acc. No, NC_045512). These were taken forward for downstream bioinformatic analyses. Here we present the analysis of prevalent lineages and the changing mutational landscape for samples collected between October 2020 to June 2022.

## Results

### Demographics and whole genome sequencing

In the study period (October 2020 to June 2022), 15,71,428 COVID-19 cases and 14,032 associated deaths were reported from the city of Bengaluru (**Figures 3a, 3b**). We started SARS-CoV-2 whole genome sequencing in August 2021 employing both retrospective and prospective genomic surveillance approach for Bengaluru. Out of 15,134 samples with collection dates ranging from October 2020 to June 2022 sequenced by amplicon sequencing approach 11,159 passed the QC of > 50% genome coverage against the reference genome. Only these genomes were considered for further analysis. Age and gender information were available for 4,424 and 4,275 (out of 11,159) samples. The median age of cases was 34 years, ranging from 1 to 92 years. In total, 2,472 were male, 1,803 were female and for 6,884 cases gender was unknown. The average age of male and female cases was 36.49 and 36.39 years, respectively (**Figure 1**).

**Figure 1:**
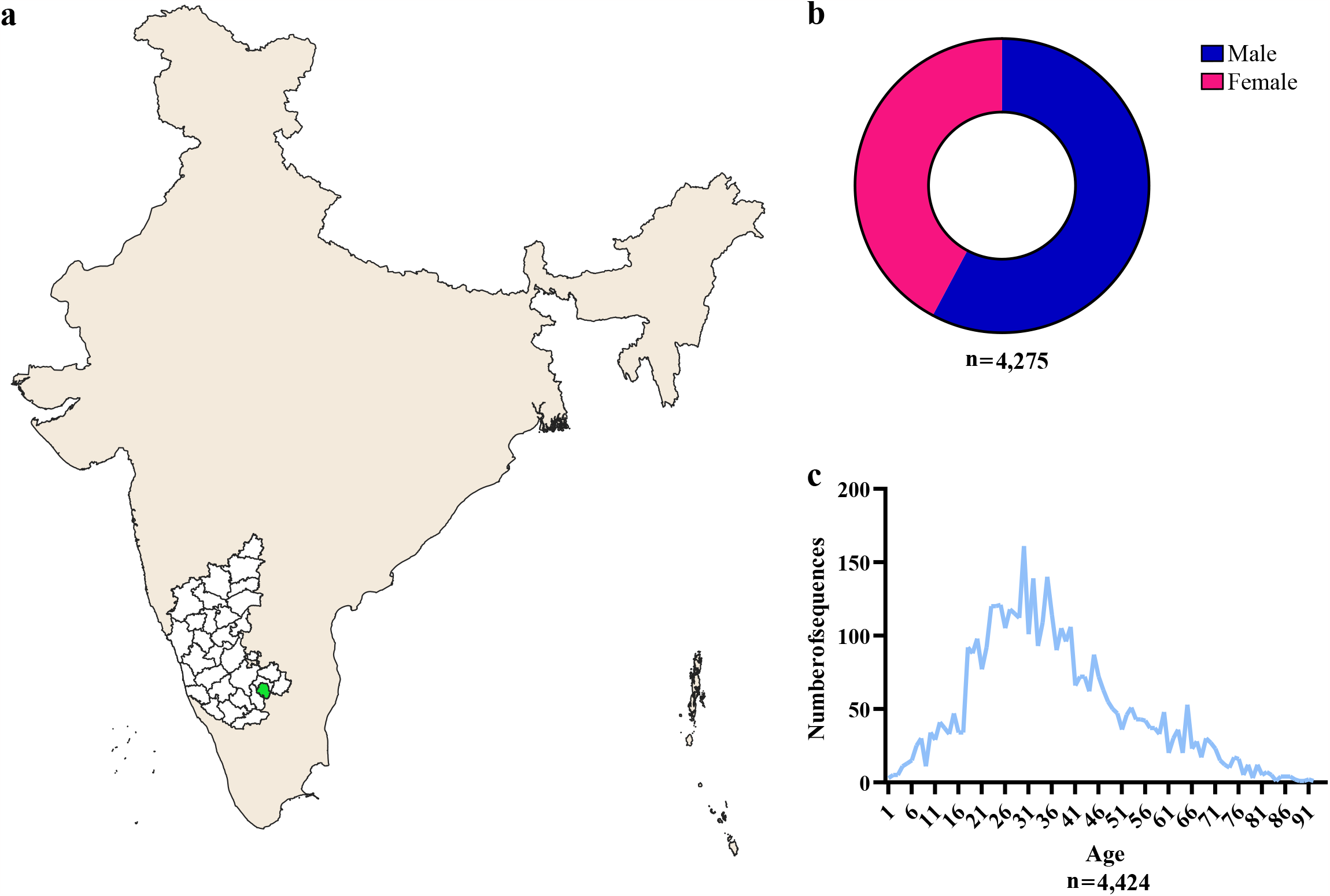
Demographics of the city level genomic surveillance in Bengaluru **a)**Geographic location of Bengaluru higlighted in green **b)**Gender distribution as parts of whole for available data in samples sequenced sequenced and passed QC **c)**Age distribution for available data in samples sequenced and passed QC

### Phylogenetic Analysis of SARS-CoV-2 Genomes

Phylogenetic analysis and lineage assignment for 11,159 assembled consensus sequences were performed using Nextclade CLI (dataset name: SARS-CoV-2-no-recomb, version 2.8.0) and PANGOLIN COVID-19 lineage assigner (version 4.1.2), respectively to identify clade and lineages. The phylogenetic analysis of 11,159 SARS-CoV-2 genomes (**Figure 2a**) revealed that the genomes were clustered into 18 major clades. A total of 196 lineages were identified by whole genome sequencing in the study. Among those, 85 lineages belonged to Delta, 77 to Omicron and 32 were other B (including Alpha, Beta, Kappa and Eta) SARS-CoV-2 lineages, respectively; the remaining were recombinant or unassigned. The detailed information of samples used in this study, such as source hospital/lab, collection date, patient age and gender, clade and lineages etc., are available in **Table S1**.

**Figure 2:**
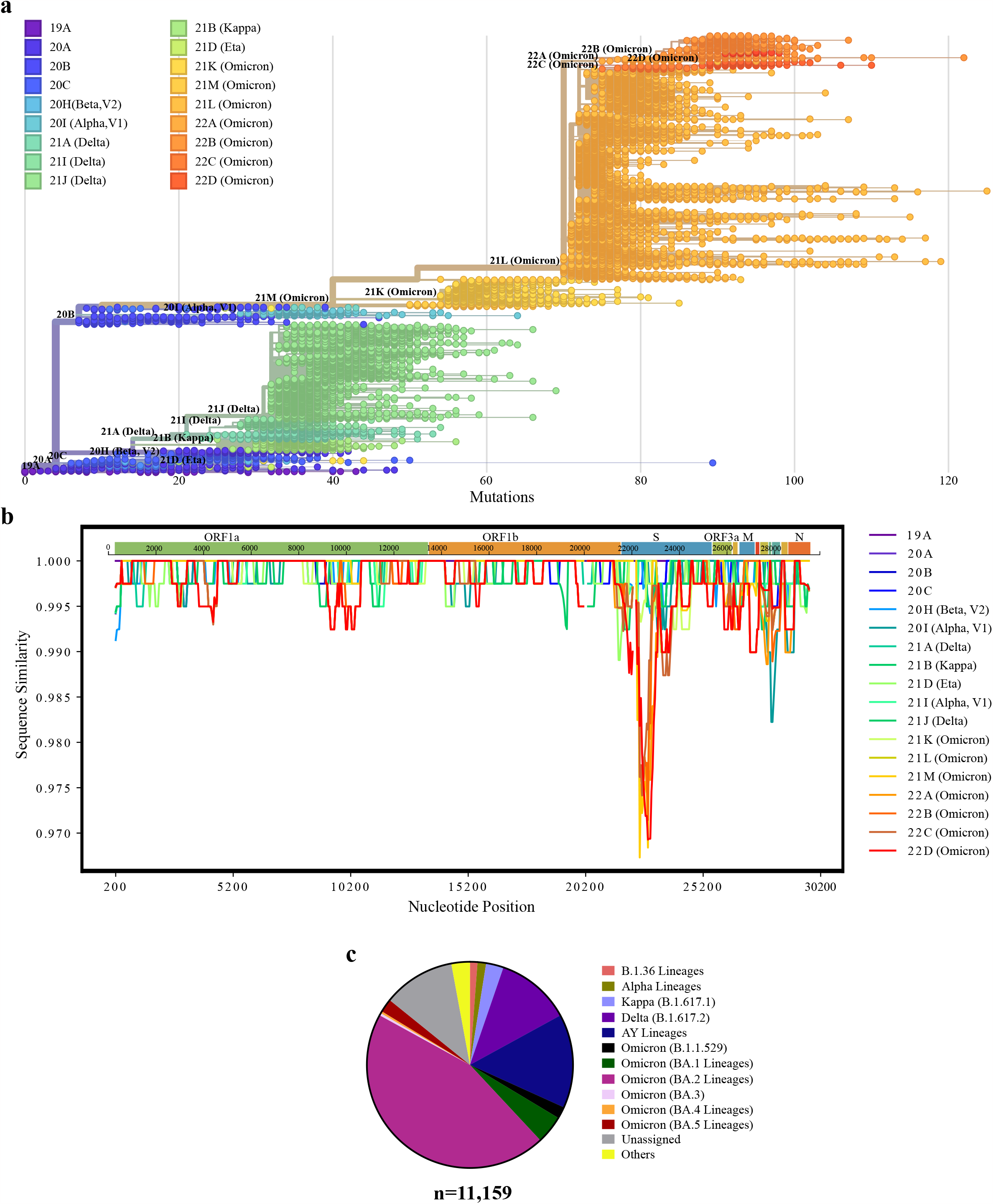
**a)** The phylogenetic analysis of 11,159 SARS-CoV-2 genomes sequenced from Bengaluru city using Nextclade showing 18 different clades. **b)** Simplot analysis showing nucleotide sequence similarity amongst different clades of SARS-CoV-2 in the study. **c)** The pie chart showing the distribution of SARS-CoV-2 lineages as parts of whole identified in this study.

### Prevalence of amino acid changes

Analysis of amino acid alterations in 11,159 SARS-CoV-2 genomes revealed a total of 7,860 unique amino acid alterations in its proteins, which includes amino acid (aa) substitutions, Insertions, and Deletions. We observed an average of 43.09 aa changes per genome. The average rate of aa substitutions, deletions and insertions were 34.97, 7.96 and 0.16 per genome respectively. Maximum number of 3,445 unique aa change events in ORF1a and a minimum of 42 unique aa changes in ORF7b protein was observed. 1,166 unique aa changes were observed in spike protein. Interestingly BA.2 lineages carried the greatest number of the aa changes. The complete list of aa changes observed from all the ORFs of SARS-CoV-2 genomes is listed in **Table S1**.

In total, 22 aa substitutions and 5 deletions were identified in ORF1a protein with above mutational prevalence cut-off and considered predominant, as mentioned in the methods. Aa substitutions S135R, T842I, G1307S, L3027F, T3090I, and the deletion of phenylalanine at the position of 3677^th^ residue was found to be more predominant in all Omicron lineages except BA.1. Whereas, A1306S, P2046L, P2287S and V2930L amino acid changes were observed only in Delta and its sublineages (AY lineages). The substitution T3255I was observed in all Omicron and Delta lineages. Further, P3395H and deletion of serine and glycine at position 3675 and 3676, respectively were predominantly observed in all Omicron lineages including the Alpha lineages. Notably, only BA.1 Omicron lineages carried few specific aa substitutions (K856R, L2084I, I3758V) and deletions (S2083-, L3674-) which were not predominant in any other lineages. T3646A was found exclusively in Kappa, Delta and its sublineages.

The aa substitutions of P314L in ORF1b protein was identified in all lineages and with dispersed prevalence. The aa substitution of K2310R in ORF1b protein was seen only in B.1.617.1 (Kappa). Moreover, the substitutions G662S, P1000L and A1918V were observed variably across the Delta lineages. Two substitutions, R1315C and T2163I showed a diverse prevalence among the Omicron lineages (except BA.1). However, I1566V was seen in all the Omicron lineages.

In the surface glycoprotein (S), 41 aa substitutions and 10 aa deletions were observed to be predominant above the mutational prevalence cut-off. T19I, S371F, T376A, D405N, and R408S mostly occurred only in Omicron lineages (except BA.1) unlike G339D, S373P, S375F, K417N, H655Y, N679K, P681H, N764K, D796Y, Q954H and N969K, which were found in all Omicron lineages. T19R, R158G, D950N and deletions E156- and F157-were found mostly in Delta and its sub lineages, whereas P681R was seen in Kappa, Delta and its sublineages. A histidine and valine deletion at position 69 and 70, respectively, were more predominant in Alpha and Omicron (BA.1, BA.4 and BA.5) lineages than in others. Another deletion (of tyrosine) at position 144 was observed mostly in Alpha and BA.1 lineages. P681H was observed in Alpha and all Omicron lineages. Further, aa changes of A67V, T95I, Y145D, T547K, N865K, L981F and deletions G142- and V143-were predominant in BA.1. Aspartic acid to glycine substitution at the position of 614 (D614G) was observed in nearly all SARS-CoV-2 lineages and it is the most frequent aa change out of all SARS-CoV-2 proteins in 95.9% (10,706 of 11,159) of this study. In addition to the changes mentioned above, other amino acid variations were also noted in the spike protein across all lineages.

The aa substitution S26L in ORF3a was found to be prevalent across Kappa, Delta and its sublineages and T223I was found in all Omicron lineages except BA.1 with varied prevalence. Interestingly in the envelope (E) protein, the mutation T9I was held only by the Omicron lineages and not by the others. The aa substitutions Q19E and A63T were observed in the membrane (M) protein of Omicron lineages, whereas I82T was identified only in Delta and its sublineages, not in others. Further, aa change from aspartic acid to glycine residue at the position of 3 in the M protein was observed more predominantly in BA.1 and less predominantly in B.1.1.529, and not observed in any other lineages.

Interestingly, aa changes in ORF7a, ORF7b and ORF8 were not found to be prevalent in any Omicron lineages but were identified in Delta and its sublineages (ORF7a: T120I, ORF7b: T40I, ORF8: D119V, F120L, D119-, F120-), with the exception of V82A in ORF7a, which was seen in Kappa, Delta and its sublineages. In ORF6, D61L was observed only in Omicron lineages of B.1.1.529, BA.2, BA.3 and BA.4 with varied prevalence.

Out of two aa substitutions observed in ORF9b, T60A was predominant in Delta and its sublineages while P10S was observed in all Omicron lineages. Three deletions in the same protein E27-, N28- and A29-were found to be prevalent in all Omicron lineages.

In the nucleocapsid protein (N), a point mutation P13L and continuous deletions of three aa residues at the position of 31 to 33 were observed primarily in Omicron lineages. Amino acid changes D63G and G215C were prominent in Delta than Omicron lineages. Amino acid substitutions R203M and D377Y were identified only in Kappa and Delta lineages but not in any other lineages (**Figure 4**).

Changing prevalence of amino acid changes across months **(Figure 5)** correlates with dominant circulating lineages in the community in respective months **(Figure 3a, 4)** and with the surge in COVID-19 cases and deaths **(Figure 5)**. Some predominant amino acid changes in Alpha lineages such as S:H69-, S:V70-, S:Y144-, S:N501Y, S:P681H, N:R203K and N:G402R that were observed to be present before the second wave were once again seen in the third wave with the emergence of Omicron lineages **(Figures 4,5)**.

**Figure 3:**
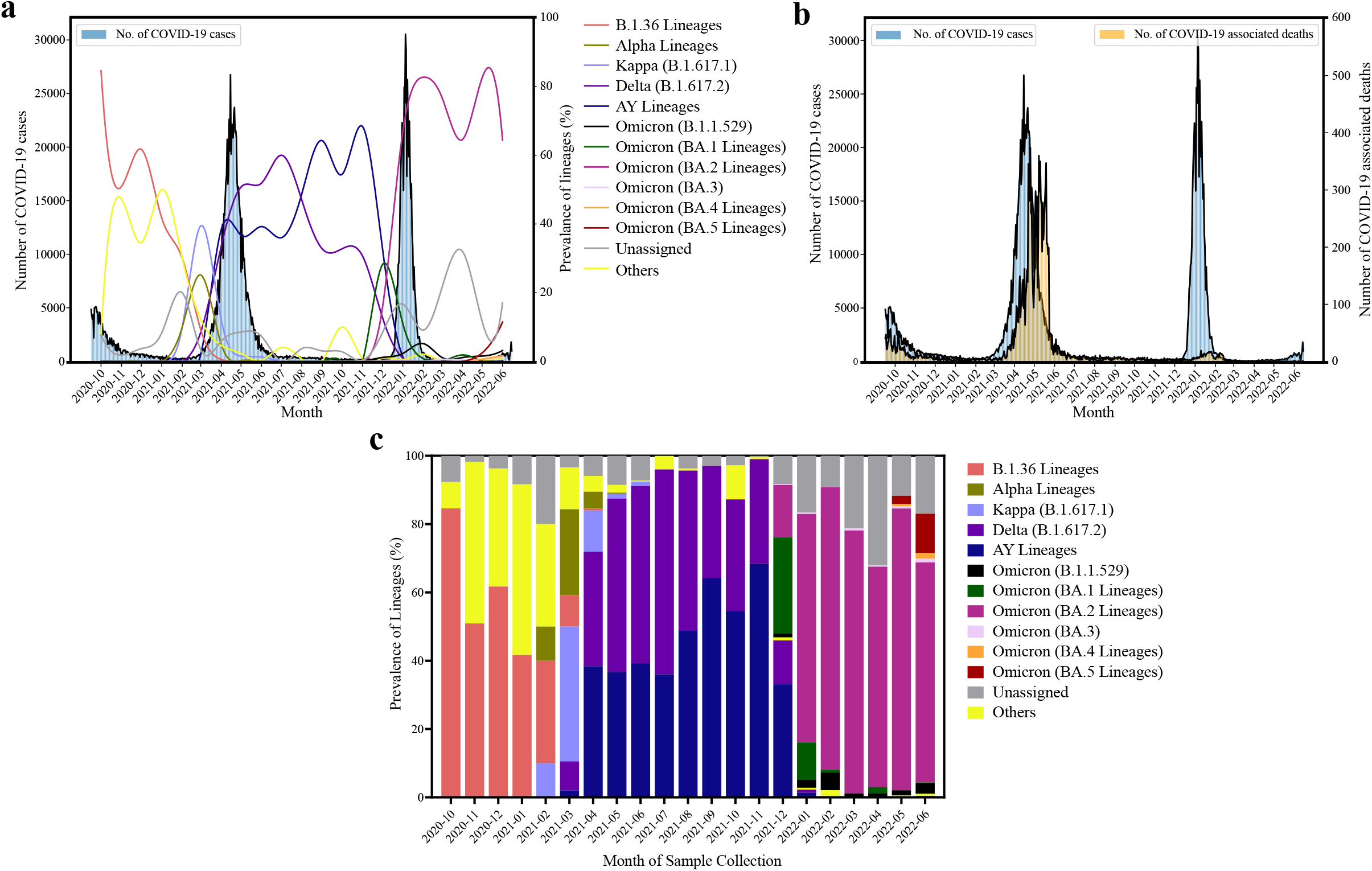
**a)** Epidemiological curve of COVID-19 cases (data obtained from https://www.incovid19.org) from October 2020 to June 2022 in the city of Bengaluru juxtaposed to proportion of SARS-CoV-2 lineages identified in the study. **b)** Representation of number of COVID-19 cases and associated deaths from October 2020 to June 2022 (data obtained from https://www.incovid19.org) in city of Bengaluru. **c)** Stacked area chart representing the distribution of SARS-CoV-2 lineages across different months in Bengaluru city (n=11,159).

**Figure 4:**
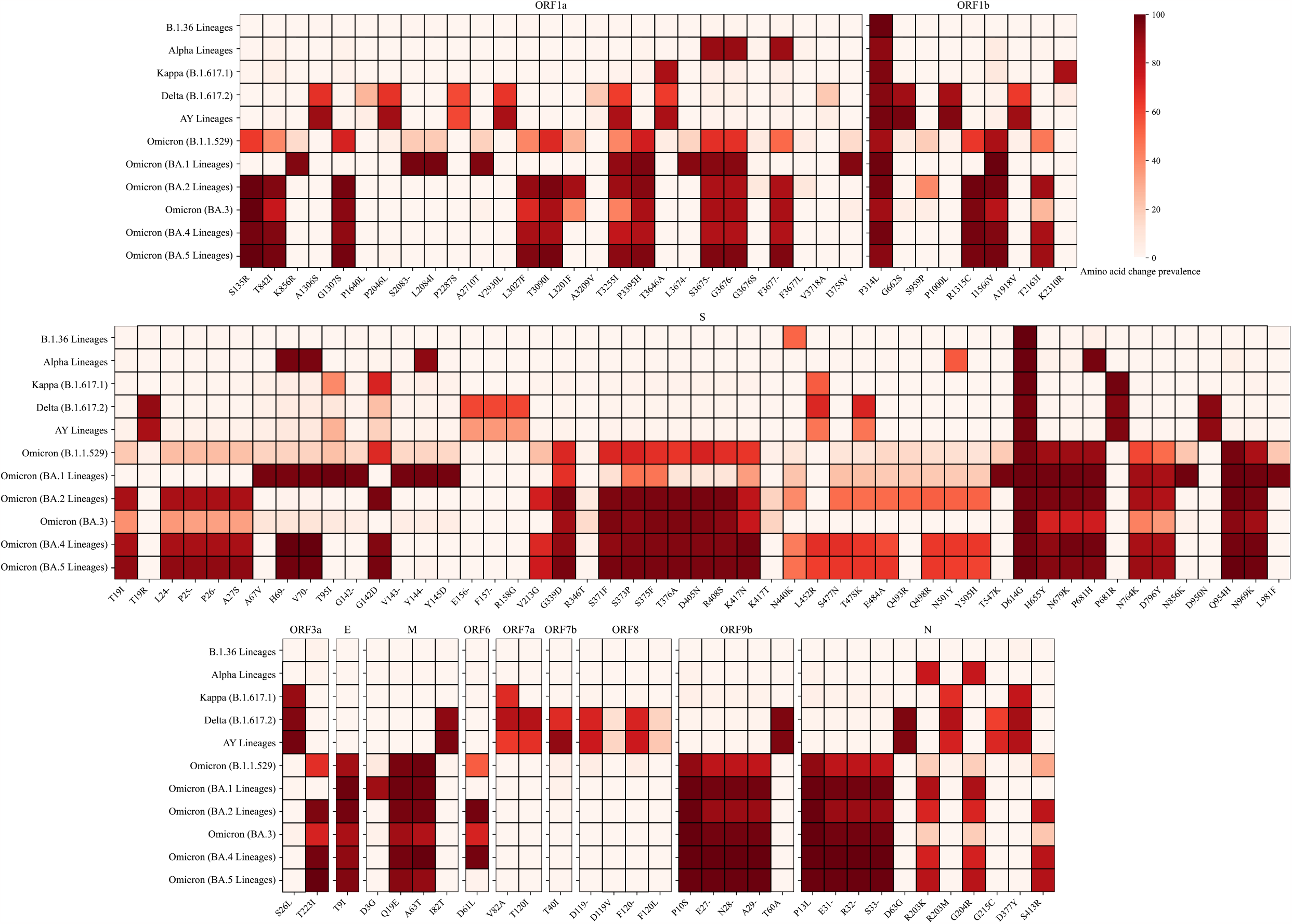
Prevalence of amino acid changes in all ORFs (names mentioned above each heatmap) of major SARS-CoV-2 lineages identified in this study.

**Figure 5:**
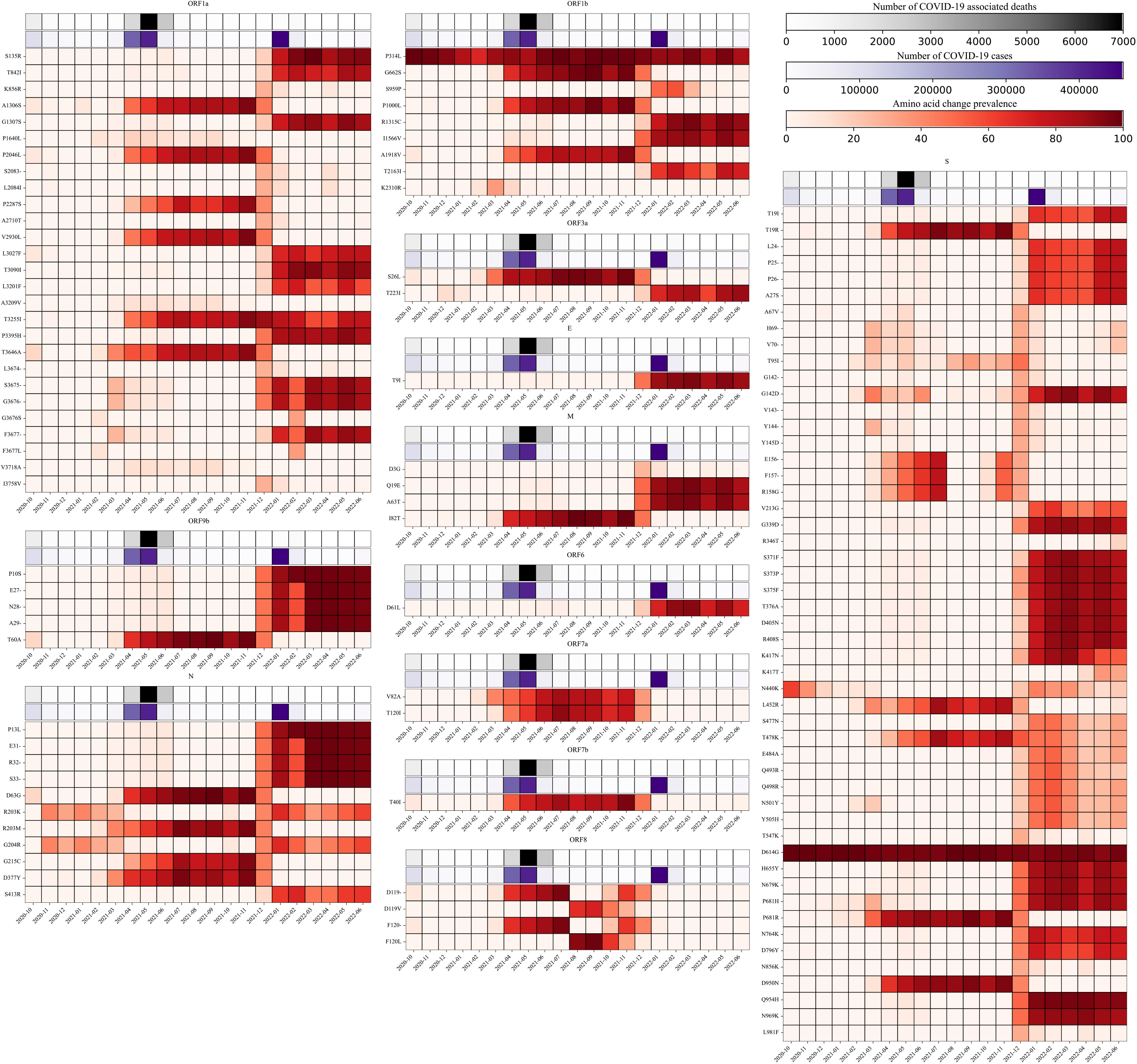
Prevalence of amino acid changes in all ORFs (names mentioned above each heatmap) of SARS-CoV-2 across months juxtaposed to number of COVID-19 cases and COVID-19 associated deaths (data obtained from https://www.incovid19.org).

### Amino acid change events landscape of spike protein among dominant lineages

The structural analysis of amino acid change events in dominant lineages in three different waves of COVID-19 cases as defined in methods reveal most of the events are observed in S1 region. At N-terminal domain of S1 more prevalent events were observed in Delta lineages followed by Omicron and least in B.1.36 lineages. Omicron lineages accounted for the greatest number of events in receptor binding domain and receptor binding motif within it. Events near S1/S2 cleavage and first heptad repeat regions were only observed in Delta and Omicron lineages not in B.1.36. All lineages were found to have events in a common region near coordinate 614 (D614G). Few events between S1/S2 cleavage and fusion peptide regions were only observed in Omicron lineages (Figure 6).

**Figure 6:**
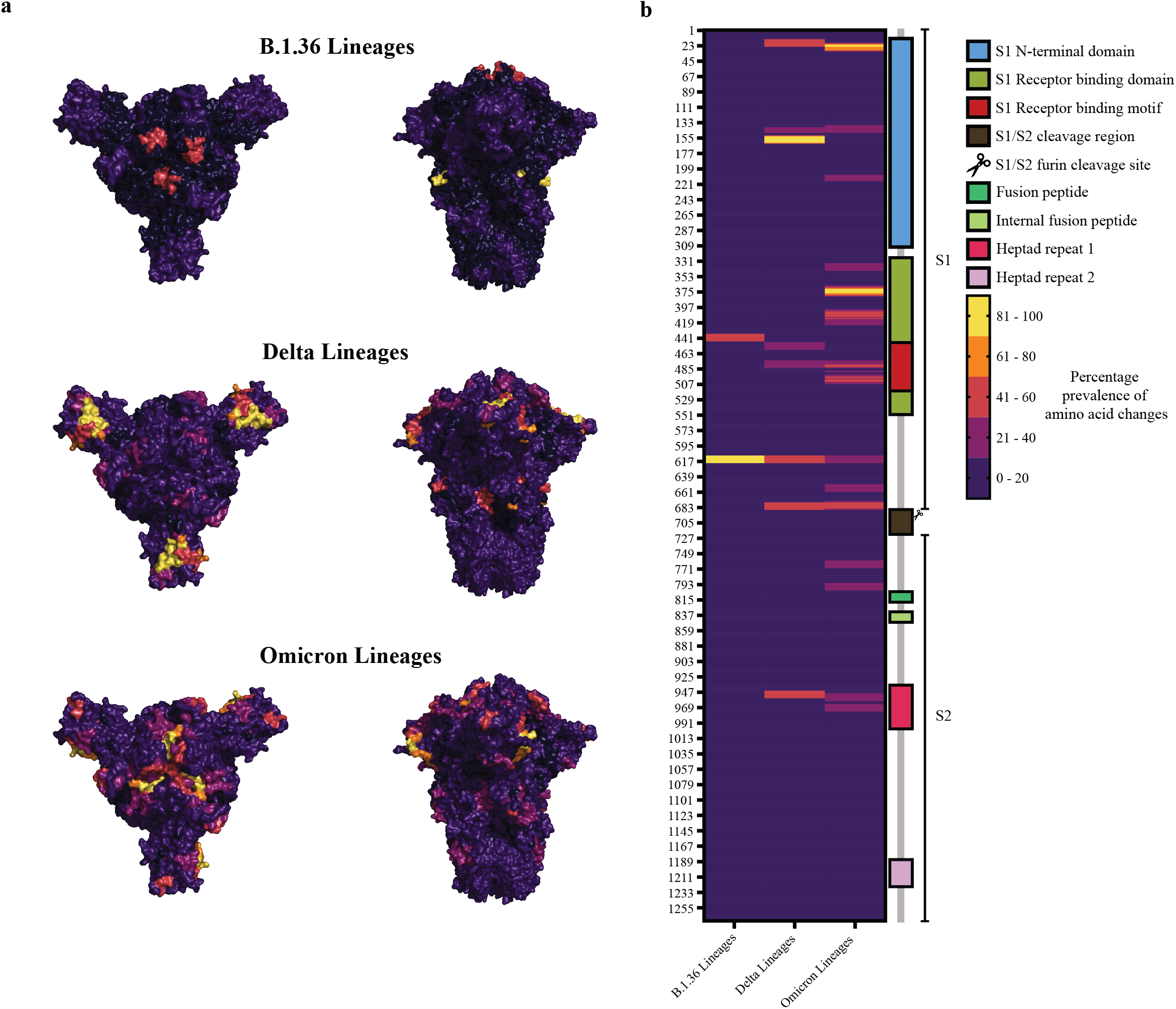
a) Prevalence of amino acid change events of B.1.36, Delta and Omicron lineages dominant in first, second, and third waves of COVID-19 mapped onto Spike protein structure (PDB 7QUS, top views on left and side views on right). b) AA change events mapped in a heatmap juxtaposed to annotation of the Spike protein.

### Distribution of amino acid changes in patient’s age and gender

The age-wise distribution of aa changes revealed that the most predominant aa changes of SARS-CoV-2 genomes with a prevalence cutoff > 3% in 11,159 sequences of this study occurred between the age of 18 to 35 years. The aa substitution D614G in spike protein was most dominant in many locales around the world and was also reported to be associated with increased infectivity^10^. This aa change in our study was most frequent in cases of the age group between 24 to 41 years **(Table S2)**. As a whole, the gender-wise distribution of aa changes revealed that the SARS-CoV-2 genomes from male cases carried higher number of aa changes (n = 3497) than female cases (n = 2736) **(Table S3)**.

## Discussion

When India encountered the first case of COVID-19 in Kerala on January 27, 2020, the Ministry of Home Affairs (MHA) announced a nationwide lockdown from March 25, 2020 to April 14, 2020 in response to increase in the COVID-19 cases across the country^9,11^. Karnataka was one of the severely affected states in India with higher number of COVID-19 cases per day during all three COVID-19 epidemic waves. Bengaluru is one of the largest city and also the capital of Karnataka with a population of around 13.1 millions. It serves as a major hub for national and international travel due to the presence of many educational institutes and universities and job opportunities, especially the globalized IT sector. The first cases of COVID-19 in the Bengaluru city were associated with international and or interstate travel^7,12^. Genomic surveillance of SARS-CoV-2 by whole genome sequencing helped in unravelling its remarkable genetic diversity and tracking of its spread worldwide. This study comprises a city level SARS-CoV-2 genomic surveillance effort by whole genome sequencing of 11,159 SARS-CoV-2 positive samples from various diagnostic labs in Bengaluru, Karnataka, India. In particular, we report the spread of SARS-CoV-2 variants with lineages and the most predominant aa changes in all the ORFs of SARS-CoV-2 genomes from Bengaluru from October 2020 to June 2022. Indeed, the SARS-CoV-2 sequence diversity in Bengaluru is remarkable and they belong to 18 clades and 196 distinct lineages.

Prior to the second wave of COVID-19 cases in Bengaluru, the B.1.36 lineages were prevalent (n=132) until the introduction of Kappa (B.1.617.1, n=307) and Alpha (B.1.1.7, a VoC, which was first detected in the United Kingdom on September 2020 with increased transmissibility and novel aa changes in the spike protein^13^, n=149). Another VoC, Beta (B.1.351) was detected in less number from March (n=2) to April 2021 (n=2). Our study was instrumental in finding Beta lineage in community; however, it was not seen to be outcompeting any other variants in circulation. Delta (AY and B.1.617.2) lineages were the most prevalent during the second wave in Bengaluru and our findings resonate with previous findings in India^7,8,12,14^. The Delta lineage was described to have originated from Indian state of Maharashtra on October 2020 and became dominant by August 2021 across the world^15^.

The B.1.1.529 lineage, was first reported from South Africa in November 2021. It was later named Omicron, and it became a dominant lineage worldwide^16^. After further evolution, B.1.1.529 got categorized into BA sublineages such as BA.1, BA.2 and so on. In our study, we detected nine cases of B.1.1.529 lineage from early December 2021. We observed that the Omicron sublineages BA.1 and BA.2 started to emerge in higher numbers in December 2021 in Bengaluru city and BA.2 became more dominant in the month of January 2022. BA.2 and its sublineages were found to be the most predominant in Bengaluru city from January 2022 to June and the trend still continued thereafter. Our study revealed that BA.2.10 (predominant between January 2022 and March 2022) and BA.2.38 (predominant between April 2022 and June 2022) sublineages accounted for most COVID-19 cases in Bengaluru city. We detected 5 and 26 cases of BA.4 and BA.5 variants respectively in Bengaluru city in late May 2022 and the numbers increased significantly [BA.4 (n = 31); BA.5 (n = 271)] by June 2022. Some prevalent aa changes associated with Alpha lineages were seen again with emergence of Omicron lineages and the phylogenetic analysis show their closer relatedness (**Figure 2a, 2c, 5, Table S1**).

The spike glycoprotein of SARS-CoV-2 is the key for its entry into the susceptible host cells. The spike protein contains the receptor binding domain (RBD) in its S1 domain which is responsible for recognition and binding to the receptor angiotensin-converting enzyme (hACE2)^17^. The RBD had been found to carry more aa changes and some of them might be crucial for increased infectivity of SARS-CoV-2^18^. The aa substitution D614G of SARS-CoV-2 spike protein had been known to be associated with enhancing entry into host cells, replication, viral load, transmission and immune system evasion^19–23^. We observed the aa substitution D614G in spike protein as most predominant with 95.9% of occurrence rate in the SARS-CoV-2 genomes. Other than D614G aa substitution in spike protein, our study demonstrated multiple lineage and sublineage specific aa substitutions and deletions in spike protein of SARS-CoV-2 genomes from Bengaluru city (**Figures 4, 5**). The same reflects in structural analysis of spike protein in all observed aa change events in all dominant lineages in three different waves of COVID-19 cases. Further, Omicron lineages dominant in third wave were found to have more aa change events compared to B.1.36 and Delta lineages dominant in the first and second waves respectively (Figure 6).

Data fetched from https://www.incovid19.org show that the greatest number of COVID-19 associated deaths were observed during second wave (**Figure 3b**) which was driven by Delta lineages and not in the third wave which also had accounted for massive numbers in daily number of COVID-19 cases. The predominant aa changes correlate with signature aa changes of dominant circulating lineages (**Figures 4, 5**). In conclusion, our study has characterized circulating variants of SARS-CoV-2 in Bengaluru city from mid of the first wave to end of third wave (October 2020 to June 2022) during the COVID-19 pandemic in India. Consistent genomic surveillance of SARS-CoV-2 circulating variants is very important for the early detection and investigation of the pattern of SARS-CoV-2 transmission. Therefore, further surveillance studies of SARS-CoV-2 are warranted to manage the COVID-19 pandemic.

## Funding Statement

This work was funded by a grant from the Rockefeller Foundation, USA to the nodal agency, CSIR-CCMB. We acknowledge support of the Department of Atomic Energy, Government of India, under project no.12-R&D-TFR-5.04-0800 and intramural funds from National Centre for Biological Sciences–Tata Institute of Fundamental Research (NCBS-TIFR).

## Data Availability

All the sequences generated out of this study and the associated metadata are available at a public domain of Global Initiative on Sharing All Influenza Data (GISAID) with GISAID Identifier: EPI_SET_221117ym, accessible at doi: 10.55876/gis8.221117ym. The data has also been deposited to CSIR Institute of Genomics & Integrative Biology (CSIR-IGIB) database.

## Acknowledgement

We are thankful to Apollo Diagnostics, Neuberg Anand Reference Laboratory, Molecular Solutions Care Health LLP, Bangalore Genomics Centre and all the laboratories for providing the samples for this study. The laboratories are acknowledged in metadata available on GISAID by Identifier: EPI_SET_221117ym, accessible at doi: 10.55876/gis8.221117ym. We thank COVID-19 testing facility at BLISC campus, Next Generation Genomics Facility at NCBS and Bhuvana Shiva for logistic help. We thank Shashidhara LS, Rakesh Mishra and Dasaradhi Palakodeti for the discussion. We also thank Bruhat Bengaluru Mahanagara Palike for the support.

## Author Contributions

DS, VK, BS, S Marate and SD collected samples from laboratories, processed samples and conducted extractions, RTPCRs, and library preparations. LC and AP supervised library preparations and sequencing. KK, DS and RN, S Mishra conducted bioinformatic analyses. UR, SM and DN designed the study. DS, KK and DN wrote the manuscript with help of all other contributing authors. DN supervised the study. All authors approved the manuscript.

## Conflict of Interest

The authors declare no conflict of interest.

## Materials and methods

### Study design and ethical consideration

For this study, nasopharyngeal and oropharyngeal swabs were collected in viral transport media (VTM) or viral lytic transport media (VLTM), that had tested positive for the presence of SARS-CoV-2 RNA by real-time reverse transcription polymerase chain reaction (RT-PCR) with cycle threshold (CT) values lesser than or equal to 28 were obtained from various diagnostic laboratories across Bengaluru. Names of the laboratories mentioned in metadata available in GISAID (doi: 10.55876/gis8.221117ym). The samples were transported to National Centre for Biological Sciences, Tata Institute of Fundamental Research (NCBS-TIFR), Bengaluru in cold chain. VTMs were processed for nucleic acid extraction in a biosafety level-2 facility at BLiSC (Bangalore Life Science Cluster) after receiving the Institutional Biosafety Committee Approval.

### Nucleic acid extraction and RT-PCR

Automated magnetic bead-based nucleic acid extractions from 200 µl VTM specimen aliquots were performed in Kingfisher flex instrument (Thermo Scientific, USA). MagMAX Dx Prefilled Viral/Pathogen Nucleic Acid Isolation kit (Invitrogen, A52076), HiPurA® Viral RNA Purification Kit (Himedia, MB615MPF-96), or MagRNA-II Viral RNA extraction kit (Genes2me, G2M030420) were used as per manufacturer’s instructions. Extracted nucleic acid was stored at -80°C until processed further.

Samples with collection dates between October 2020 to July 2021 retrieved from biorepositories as part of our retrospective study were subjected to RT-PCR screening using CoviDxT mPlex-4R kit (NeoDx, CM4R01). Retrospective RT-PCR positive samples with CT values lesser than or equal to 30 were further processed for sequencing.

### Library Preparation and Whole Genome Sequencing

The libraries were prepared using Illumina COVIDSeq Test (RUO) kit (Illumina, 20043675) as per manufacturer’s instructions at the Next Generation Genomics Facility of NCBS-TIFR, Bangalore. The preparations for first strand and PCR amplification of SARS-CoV-2 whole genome with specific primer sets targeting SARS-CoV-2 RNA along with 11 primers targeting human RNA serving as control were carried out in a BSL-2 hood. Amplicons were tagmented using enriched bead linked transposase, indexed using IDT for Illumina Nextera UD indexes Sets 1–4 (384 Indexes, Cat no: 20043137, Illumina, USA) and products were amplified. Libraries were uniformly pooled and purified using Illumina Tune Beads (0.9x). The pooled libraries were quantified using Qubit 4.0 Fluorometer (Invitrogen, USA) and library sizes were analyzed on TapeStation 4200 (Agilent, USA). The pooled libraries were further normalized to 2nM concentration and were denatured using 0.1N sodium hydroxide. 8.1pM of denatured libraries were loaded onto HiSeq Rapid SR flow cell v2. Dual indexed (10bp) 50, custom 100 or 120 cycles single read runs were performed on HiSeq 2500 instrument (Illumina, USA) using HiSeq Rapid SBS kit v2 (50 cycle) kit. The raw sequencing data in binary base call (BCL) format were converted into fastq using bcl2fastq version2.20, after de-multiplexing.

### Genome assembly, Phylogeny and Lineage assignment

The raw reads were demultiplexed to FASTQ using bcl2fastq (version 1.8) and uploaded to Illumina BaseSpace via its command line interface and subjected for genome assembly using DRAGEN COVID Lineage app (version 3.5.3, https://basespace.illumina.com/apps/12139127/DRAGEN-COVID-Lineage). 11,159 consensus sequences generated with >50% coverage by mapping against reference genome were shared on GISAID. Data for lineage assignment using Phylogenetic Assignment of Named Global Outbreak LINeages (PANGOLIN, version 4.1.2, consensus calls)^24^, was retrieved from GISAID^24^. The phylogenetic tree was constructed using Nextclade command line interface (dataset name: SARS-CoV-2-no-recomb, version 2.8.0) with assembled genomes against the global reference dataset (genbank acc. No, NC_045512) and visualized using auspice interactive phylogenetic tree visualizing tool from Nextstrain (**Figure 2a)**^25^.

### Simplot Analysis

The consensus nucleotide sequence similarity of different clades observed in our study to the reference sequence of SARS-CoV-2 was estimated for every 400bp with step size 50 bp using Kimura’s two nucleotide substitution model of evolution (**Figure 2b**)^26^.

### Statistical analysis

All data were represented as counts and percentages. The aa changes with more than 3% prevalence in 11,159 sequences in this study were plotted in heatmaps for their distribution across group of lineages (**Table S1**) and month of sample collection (**Figures 4, 5**). Daily number of COVID-19 cases and associated deaths were obtained from https://www.incovid19.org (**Figures 3a,3b**).

### Structural mapping of Spike protein amino acid change events in dominant lineages across COVID-19 waves

Sequences of B.1.36 (n=89, sample collection dates between October 2020 and December 2020), Delta (n=1765, sample collection dates between March 2021 and July 2021), and Omicron (n=3344, sample collection dates between December 2021 and February 2022) (sub)lineages which were dominant in first, second and third waves of COVID-19 cases were included for mapping of all aa change events on publicly available Spike protein trimer structure (PDB 7QUS) using PyMOL^27^. Rolling sum of aa change events in a 5 aa flank was mapped to each residue in the linear protein sequence, which were subsequently min max normalized and split into quantiles. The quantiles were then mapped onto the Spike protein structure.

## Notes

### Competing Interest Statement

The authors have declared no competing interest.

